# Host-plant adaptation as a driver of incipient speciation in the fall armyworm (*Spodoptera frugiperda*)

**DOI:** 10.1101/2022.09.30.510290

**Authors:** Estelle Fiteni, Karine Durand, Sylvie Gimenez, Robert L. Meagher, Fabrice Legeai, Gael J. Kergoat, Nicolas Nègre, Emmanuelle d’Alençon, Kiwoong Nam

## Abstract

**Background:** Divergent selection on host-plants is one of the main evolutionary forces driving ecological speciation in phytophagous insects. The ecological speciation might be challenging in the presence of gene flow and assortative mating because the direction of divergence is not necessarily the same between ecological selection (through host-plant adaptation) and assortative mating. The fall armyworm (FAW), a major lepidopteran pest species, is composed of two sympatric strains, corn and rice strains, named after two of their preferred host-plants. These two strains have been hypothesized to undergo incipient speciation, based on (*i*) several lines of evidence encompassing both pre- and post-zygotic reproductive isolation, and (*ii*) the presence of a substantial level of genetic differentiation. Even though the status of these two strains has been established a long time ago, it is still yet to be found whether these two strains indeed exhibit a marked level of genetic differentiation from a large number of genomic loci. Here, we analyzed whole genome sequences from 56 FAW individuals either collected from pasture grasses (a part of the favored host range of the rice strain) or corn to assess the role of host-plant adaptation in incipient speciation.

**Results:** Principal component analysis of whole genome data shows that the pattern of divergence in the fall armyworm is predominantly explained by the genetic differentiation associated with host-plants. The level of genetic differentiation between corn and rice strains is particularly marked in the Z chromosome. We identified one autosomal locus and two Z chromosome loci targeted by selective sweeps specific to rice strain and corn strain, respectively. The autosomal locus has both increased D_XY_ and F_ST_ while the Z chromosome loci had decreased D_XY_ and increased F_ST_.

**Conclusion:** These results show that the FAW population structure is dominated by the genetic differentiation between corn and rice strains. This differentiation involves divergent selection targeting at least three loci, which include a locus potentially causing reproductive isolation. Taken together, these results suggest the evolutionary scenario that host-plant speciation is a driver of incipient speciation in the fall armyworm.

## INTRODUCTION

Host-plant adaptation is one of the main evolutionary forces causing ecological speciation in phytophagous insects [1] since plants provide nutrients, oviposition sites, and mating places. Population genomics and molecular evolutionary analyses provide powerful tools to identify adaptively evolved insect genes potentially causing host-plant adaptation. These genes encode chemosensory proteins to detect suitable plants, oral secretion proteins to respond to plant defense, digestion genes to catabolize plant molecules, and detoxifying proteins to neutralize plant secondary metabolites [2, 3]. Several studies also show that these genes exhibit accelerated adaptive evolutionary rates in phytophagous insects [4–6]. Interestingly, polyphagous phytophagous insects generally have higher numbers of detoxification and chemosensory genes than monophagous ones [7–10] probably due to the consequence of the interactions with diverse plant molecules from diverse plant species [11].

In the presence of gene flow, speciation by host-plant adaptation can be challenging. Typical speciation processes with gene flow involve both prezygotic reproductive isolation by assortative mating and postzygotic reproductive isolation by ecological divergent selection (such as divergent selection on the usage of host-plants) [12]. As demonstrated by the classical paper by Felsenstein [13], recombination between genetic loci determining assortative mating and ecological divergent selection generates all allelic combinations for these loci, and evolutionary trajectories of divergence are determined by the relative strength between ecological divergent selection and assortative mating. Therefore, the presence of divergent selection on host-plants does not necessarily imply that speciation will occur between two populations with different host-plants. Since the 1990s, theoretical evolutionary studies have shown that speciation may occur even in the presence of gene flow in particular sets of conditions overcoming the homogenizing effect of recombination, and almost a hundred models of speciation with gene flow have been proposed[12].

For example, if host-plants provide both nutrients and mating sites, such as in the case of the *Rhagoletis pomonella* sibling-species complex [14, 15], recombination does not affect divergence because there is only one trait causing the divergence [16, 17]. In this case, speciation may occur readily between a pair of sympatric populations.

The fall armyworm (FAW), *Spodoptera frugiperda* (Lepidoptera: Noctuidae: Noctuinae) is a major pest species native to the Americas that recently invaded the Eastern hemisphere, with invasive populations being first reported in West Africa in 2016 [18]. Since then, it quickly spread in almost all of sub-Saharan Africa, and then progressively expanded its range in Egypt, Asia, and Australasia (https://www.cabi.org/isc/fallarmyworm), and the FAW is considered one of the worst invasive pest species in Africa [19]. The FAW consists of two ecologically divergent host-plant strains, referred to as the corn strain (sfC) and rice strain (sfR) [20, 21]. Even though the FAW is a very opportunistic and polyphagous pest species [22], sfC and sfR strains are known for displaying differentiated ranges of preferred host-plants, such as sfC prefers corn, sorghum, and cotton, whereas sfR prefers rice, millet, and pasture grasses. The two strains are observed in sympatry in the FAW native range. Hybrid individuals have been also documented with proportions as high as 16% [23]. Reciprocal transplant experiments demonstrated that the two strains present differential performances on their preferred host-plants [24], which implies the existence of differential hostplant adaptation. Interestingly, sfC and sfR have allochronic mating patterns [25, 26] and different compositions of sexual pheromone blends [27, 28], and hybrid crosses generated in a lab have reduced fertility [29], implying a possibility that host-plant adaptation might not be a single evolutionary force causing divergence in FAWs. The status of both strains has been often questioned [30, 31], and the extant consensus is that these two strains are engaged in a process of incipient speciation [32]. Mitochondrial cytochrome *c* oxidase subunit 1 (COX1) gene [33, 34] and Z chromosome Triosephosphate isomerase (TPI) gene [35] have been widely used to identify both strains.

Several studies demonstrated that genomic differentiation occurs between sfC and sfR strains. For example, Tessnow et al. [36] showed from samples collected from Texas that sfC and sfR have allochronic matings as well as genomic differentiation. Durand et al. [37] analyzed whole genome resequencing data, originally generated by Schlum et al. [38], of 55 samples collected from Argentina, Brazil, Kenya, Puerto Rico, and the mainland USA. They also observed that whole genome sequences are differentiated between sfC and sfR samples, partly due to very strong divergent selection on Z chromosomes, which caused autosomal differentiation by genome hitchhiking [39]. It should be noted that most samples in these studies were collected at the adult stages near corn or sorghum, which are known to belong to the preferred host-plants of sfC. Therefore, the effect of host-plant during larval stages in the incipient speciation of FAW is still unknown. We first reported genomic differentiation between strains from larval samples collected from a corn field in Mississippi [8, 40]. However, larval samples from sfR-preferred host-plants were not included in these studies. In short, the effect of host-plant in genomic differentiation is yet to be reported.

In this study, we analyzed whole genome sequences of FAW samples at the larval stage that were collected from corn fields (one of the sfC preferred host-plants) and a pasture grass field (part of the sfR preferred host range) to test whether differential host-plant adaptation drives incipient speciation between sfC and sfR. First, we test whether the population structure of FAW is mainly determined by the differential ranges of host-plants. Second, we test the existence of divergent selection that potentially caused differential adaptation to host-plants. Third, we test the genetic differentiation of the *vrille* gene, which was shown to determine allochronic mating patterns in FAW [25]. It should be noted that we do not test for the possibility that speciation occurs only through differential host-plant adaptation. Instead, we aim at testing the major effect of host-plant adaptation during potential incipient speciation in the FAW.

## RESULTS

### Reference genome assembly, strains, and resequencing dataset

The size of the assembled reference genome was 385 Mb, and N50 is 10.6 Mbp. L90 is 26, which is close to the known number of chromosomes in the FAW (31), implying that we nearly have chromosome-sized scaffolds in this assembly. The BUSCO analysis [41] showed that this assembly had the highest correctness among all published FAW genome assemblies (Table S1). The number of identified SNVs (single nucleotide variations) from 56 samples was 22,877,074.

When the TPI locus was used to identify strains, an almost perfect correlation between host-plants and the identified strain was observed (Table 1, Table S2, and Fig. S1), with the single exception that an sfR individual was found from a corn field (MS_R_R6). When the mitochondrial COX1 was used, one and ten samples from the pasture grass were assigned to sfC and sfR, respectively. The numbers of sfC and sfR samples from corn fields were 33 and 12, respectively.

**Table 1.** The numbers of identified sfC and sfR samples using the TPI or the mitochondrial COX1 markers.

| host-plants | TPI |  | mtCOX1 |  |
| --- | --- | --- | --- | --- |
|  | sfC | sfR | sfC | sfR |
| Corn field | 44 | 1 | 33 | 12 |
| pasture grass | 0 | 11 | 1 | 10 |

### The effect of host-plants on genetic differentiation

Principal component analysis of whole genome data recovered two groups at the first principal component (Fig. 1A). This grouping had a perfect correlation with host-plants with a single exception of a single sample from corn (MS_R_R6), which was clustered with other samples from pasture grasses. Here, we categorized the groups composed of samples from corn and pasture grasses as the corn group and the grass group, respectively. All the samples from the corn group and the grass group can be assigned to sfC and sfR according to the TPI marker, respectively. All samples from the grass group were assigned to sfR identified based on the mitochondrial COX1 marker except for one sample (FGJ4). In the corn group, 33 and 11 samples were assigned to sfC and sfR according to the mitochondrial COX1 marker, respectively. Interestingly, all 13 samples from the corn group in Florida were sfC according to the mitochondrial COX1 marker, whereas only 62.5% of the corn group in Puerto Rico and Mississippi were sfR (20 out of 32). The grouping according to geographic population was not observed within the corn group from the first to tenth principal components (Fig. S2).

**Figure 1.**
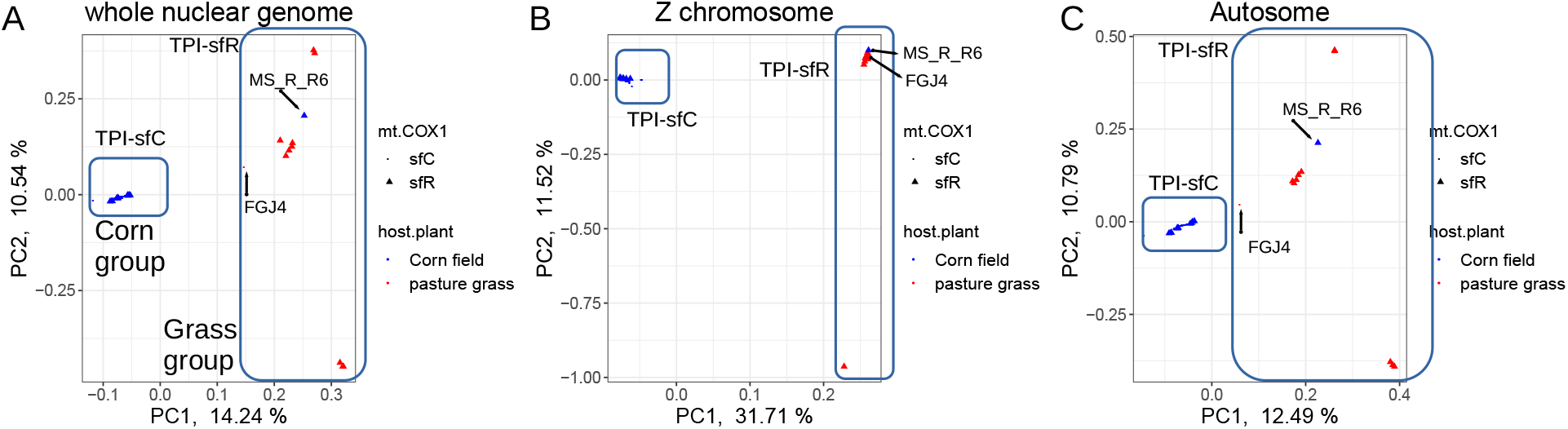
Genetic differentiation between host-plants and strains. Principal component analysis from (A) whole nuclear genome, (B) the Z chromosome, and (C) autosomes. TPI-sfC and TPI-sfR represent strains identified from the TPI marker. The red and blue points indicate samples collected from corn fields and pasture grasses, respectively.

When the principal component analysis (PCA) was performed from the Z chromosome and autosomes separately, the same trend was observed (Fig. 1B and C). Notably, the Z chromosome PCA showed that FGJ4 was found within the corn group, whereas the autosomal PCA results indicated that FGJ4 was closest to the grass group along with the first principal component. This result implied the possibility that FGJ4 is a hybrid between sfC and sfR. Therefore, we performed ancestry coefficient analysis to test this possibility from the samples from Florida. sfC and sfR samples exhibited differentiated ancestry, while FGJ4 had almost the same proportions of ancestry between sfC and sfR (Fig. S3).

We further tested genetic differentiation using FST statistics. The average FST between the corn group and the grass group was 0.0813. To test whether this F_ST_ value can be generated by chance, F_ST_ was calculated from random grouping with 100 replications. All 100 replications had F_ST_ lower than 0.0813 (equivalent to *p-value* < 0.01) (Fig. 2A). F_ST_ from the Z chromosome was 0.4603, which is far higher than all autosomal chromosomes, as shown from previous studies [36, 37]. In total, 100% of untruncated 500kb windows have F_ST_ higher than zero and statistically significant genetic differentiation was observed from 99.6% of the windows (FDR corrected p-value < 0.05). These results imply genomic differentiation (GD), which was defined as a status where genetic differentiation occurred in a vast majority of loci (e.g., > 90%) across the whole genome [40].

**Figure 2.**
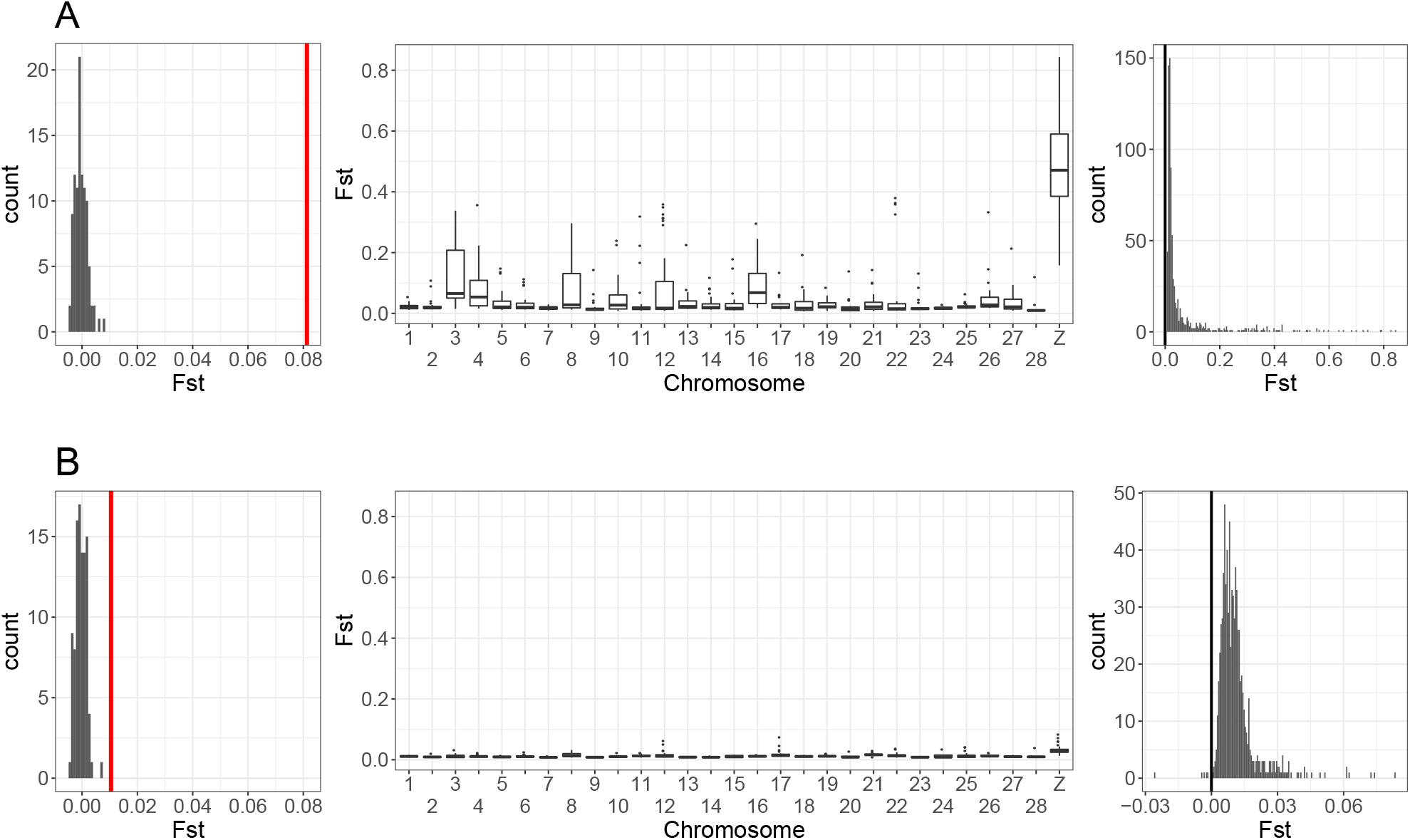
Genetic differentiation between groups in the FAW. A. FST was calculated between the corn group and the grass group. (left) The histogram shows FST calculated from random groups. The red vertical bar indicates FST calculated between the corn group and the grass group. (middle) FST calculated in 500kb windows was shown for each chromosome. (right) The histogram of F_ST_ was calculated in 500kb windows. The black vertical bar indicates F_ST_= 0. B. F_ST_ was calculated between two groups with different mitochondrial markers within the corn group. Please note that A and B have different ranges of FST in the rightmost panels.

F_ST_ between sfC and sfR from the corn field was only 0.0105. However, none of the 100 replications with random grouping had F_ST_ lower than 0.0105 (Fig. 2B), which implies a statistically significant genetic differentiation (*p-value* <0.01), as shown previously [8, 40, 42]. F_ST_ calculated from the Z chromosome was 0.0292, which was slightly higher than autosomes. We observed that 99.60% of untruncated 500kb windows have F_ST_ higher than 0, and 92.7% of these windows exhibited statistically significant genetic differentiation. These results imply GD between sfC and sfR within the corn group.

### Divergent selection between corn and grass groups

Targets of divergent selection were identified from genetic footprints of selective sweeps using the composite likelihood approach[43]. We considered outliers of composite likelihood specific to the corn or grass groups to be targets of selective sweeps to minimize the possibility of background selection[44, 45]. The grass group had one obvious outlier reflecting the composite likelihood of selective sweep on chromosome 12, while the corn group had two outliers on the Z chromosome (Fig. 3, Fig. S4, and Table S3). Four genes were identified from the grass group-specific outlier, but the function of these genes is unclear (Table S4). The two corn group-specific outliers had 58 genes, which include 47 genes with unknown functions.

**Figure 3.**
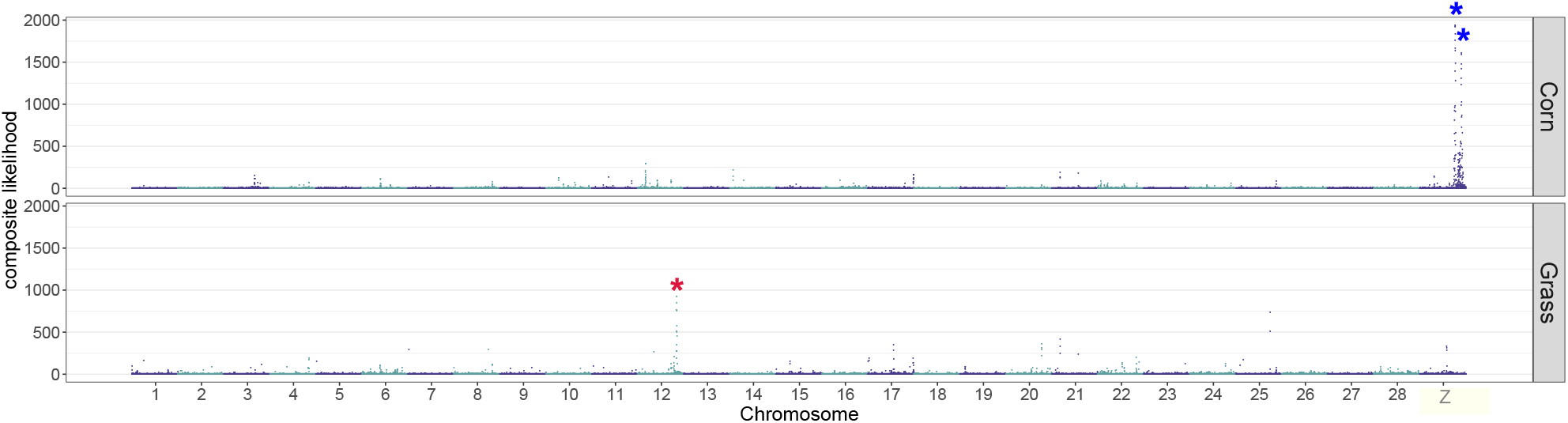
Selectively targeted loci. The composite likelihood of selective sweep along the genome was calculated from the corn or grass group. Obvious outliers of likelihood were indicated by asterisks

If a selectively targeted locus caused reproductive isolation, then this locus is expected to have an increased level of absolute differentiation (i.e., D_XY_) because the reduced rate of gene flow causes an ancient divergence time, and an increased level of relative differentiation (i.e., F_ST_) because natural selection removes shared SNPs between populations[46]. Our forward simulation showed that divergently targeted loci with reduced gene flow exhibited increased FST and DXY (Fig. S5), confirming this expectation. Then, we calculated DXY and FST from the chromosomes containing the identified targets of selective sweeps. The grass group-specific outlier on chromosome 12 had increased DXY and increased FST (Fig. 4A). The two corn group-specific outliers on the Z chromosome showed increased FST, but increased DXY was not observed from these outliers (Fig. 4B).

**Figure 4.**
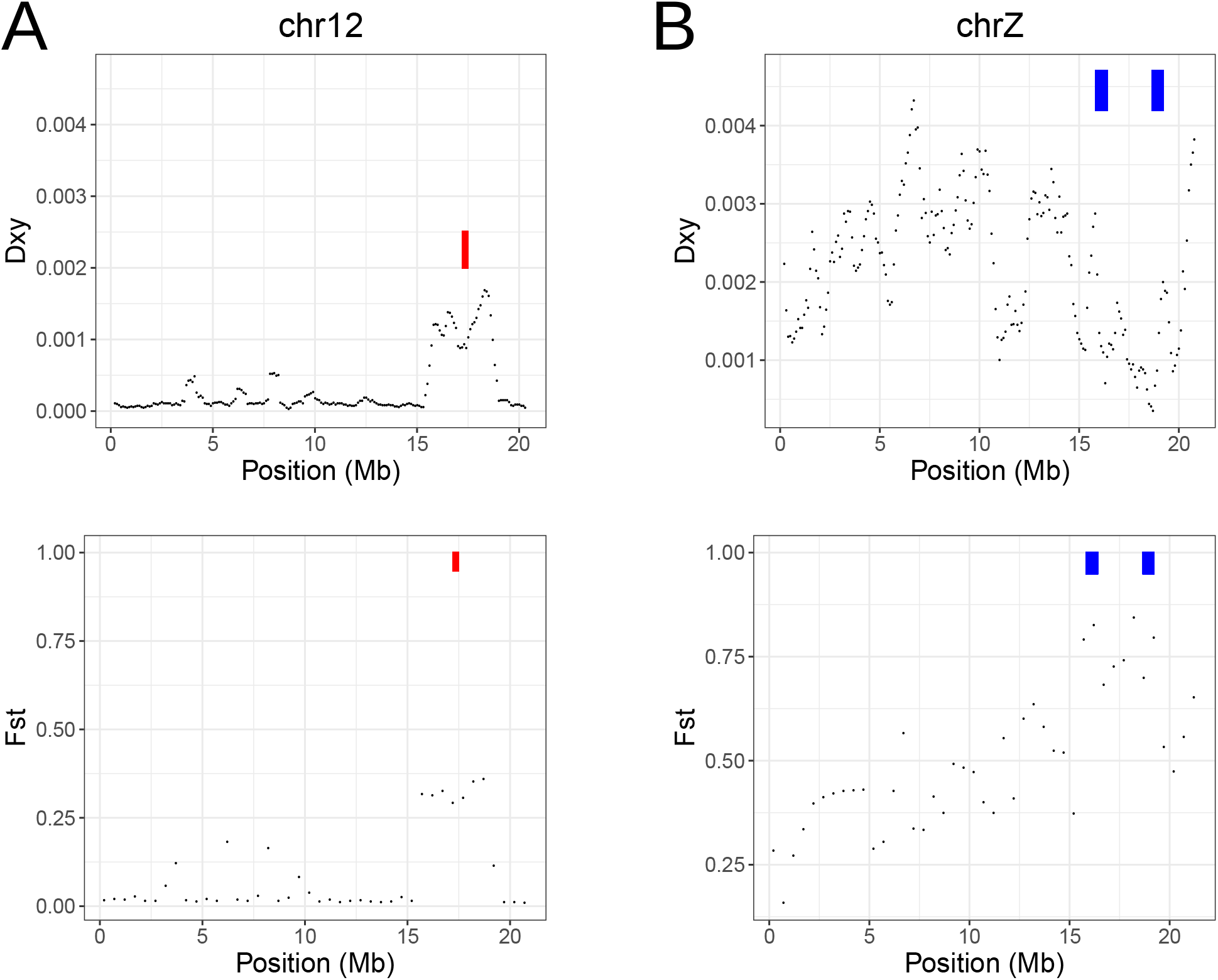
Loci under divergent selective sweeps. D_XY_ (upper) and F_ST_ (lower) were calculated from the targets of selective sweep specific to the grass group (the red rectangles) and the corn group (the blue rectangles).

The *vrille* gene was reported to cause allochronic mating behavior in FAW through QTL mapping [25]. Thus, we tested whether there is an elevated genetic differentiation at the *vrille* gene. Interestingly, FST does not appear to be particularly higher at the *vrille* gene than the chromosomal average (Fig. S6).

## DISCUSSION

Divergent selection on host-plants is often considered to be one of the main evolutionary forces driving speciation in phytophagous insects. In this study, we showed that the FAW is composed of two genomically differentiated groups with different host-plants, the corn group and the grass group, based on population genomics analyses (Fig. 1 and Fig. 2A). The ancestry coefficient analysis supported the existence of hybrids (FGJ4), suggesting the presence of gene flow (Fig. S3). We identified three loci that were targeted by corn or grass group-specific selective sweeps (Fig. 3), suggesting the possibility that divergent selection contributed to the genetic differentiation between the corn and the grass groups. The grass group-specific target had both increased D_XY_ and F_ST_ (Fig. 4A), implying that divergent selection on this locus caused reproductive isolation. Intriguingly, the two corn group-specific targets did not have increased DXY (Fig. 4B), making the link between divergent selection and reproductive isolation unclear. Taken together, we conclude that the FAWs analyzed in this study are composed of two genomically differentiated groups with differentiated ranges of host-plants and that divergent selection contributed to the speciation process between these two groups. Interestingly, we also observed genetic differentiation between the two mitochondrial strains within the corn group (Fig. 2B).

We propose the following evolutionary scenario of speciation in FAW from these results (Fig. 5).*i*)Divergent selection targeting chromosome 12 caused reproductive isolation between ancestral corn and grass groups (Fig. 4A). The ancestral corn group experienced divergent selection on the Z chromosome (Fig. 4B). As a consequence, extant corn and grass groups had differentiated ranges of host-plants with differentiated genomic sequences (Fig. 1 and Fig. 2A).*ii*) Following evolutionary forces caused the nuclear divergence of the corn group into two sub-groups (Fig. 2B), possibly involving mild divergent selection targeting many loci [40]. These two sub-groups had different mitochondrial genomic sequences including the COX1 genes (Fig. S1) for a reason yet to be identified. We suggest that the corn group and the grass group should be considered as sfC and sfR, respectively. Here, the two sub-groups within the corn group can be presumably named mt-A and mt-B, rather than sfC or sfR.

**Figure 5.**
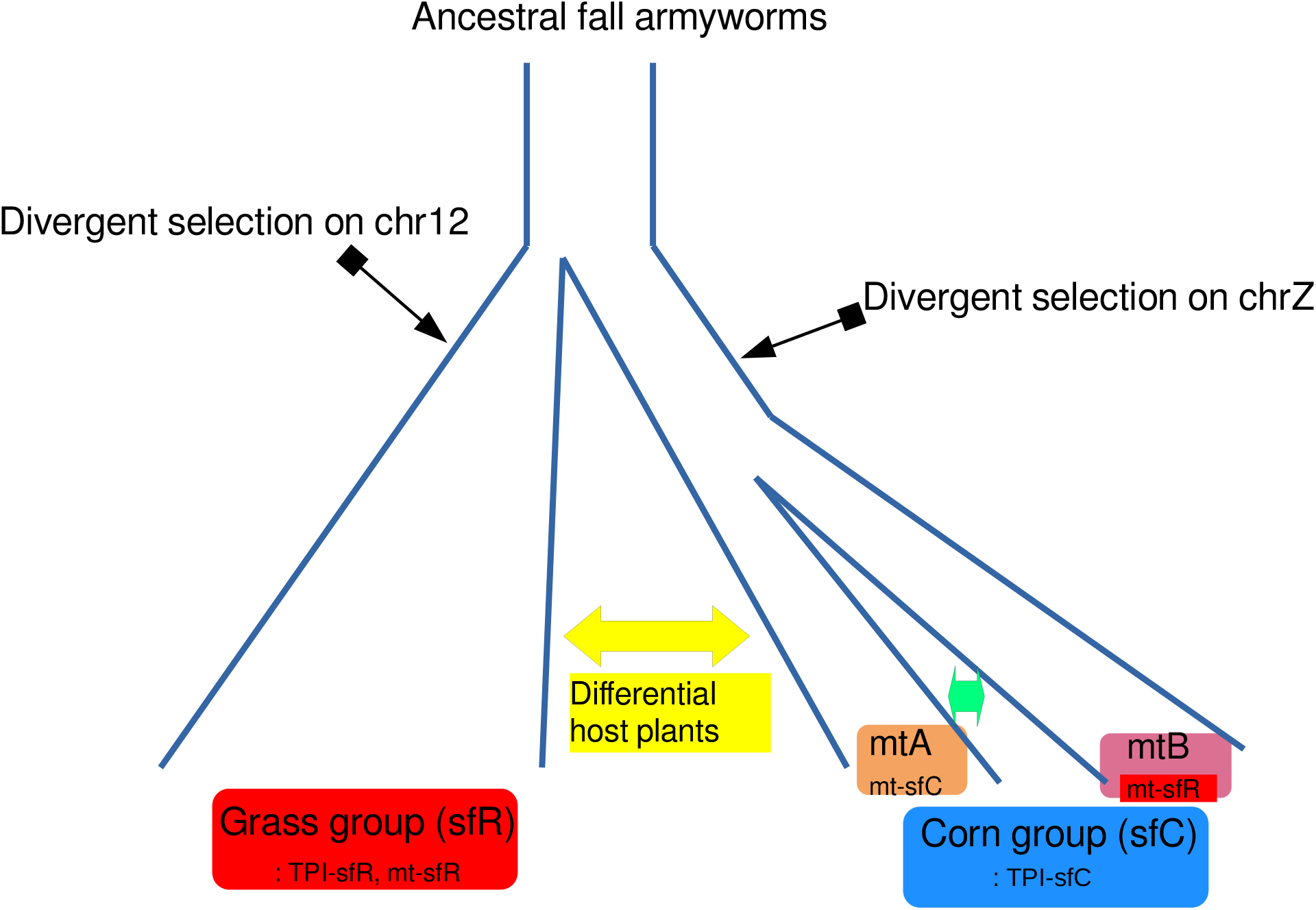
An evolutionary scenario of speciation in the fall armyworm. *i*) Divergence selection on a locus on chromosome 12 caused reproductive isolation by reducing gene flow between ancestral corn and grass groups. The ancestral corn group experienced divergent selection on the Z chromosome as well. As a consequence, the corn group (sfC) and the grass group (sfR) had differentiated ranges of host-plants. *ii*) Evolutionary forces split the corn group (sfC) into two subgroups, mt-A and mt-B, with different mitochondrial markers.

Interestingly, genetic differentiation was observed across almost the entire genomic loci (i.e., GD), between sfC (the corn group) and sfR (the grass group), and between mt-A and mt-B. In other words, the significant genomic differentiation between sfC and sfR or between mt-A and mt-B is caused by very large numbers of loci with low genetic differentiation, rather than small numbers of loci with high genetic differentiation. Geographic separation is not likely to be a plausible explanation because the strong migratory behavior of FAW [47] likely causes genetic admixtures between the two geographic populations within ~150 km distance (i.e. the grass group and the corn group in Florida), through gene flow. In the presence of such gene flow, only loci targeted by divergent selection are expected to be differentiated between intraspecific races [48]. Moreover, mt-A and mt-B were collected from the same field. Indeed, we did not observe population structure according to the geographic populations within the analyzed sfC samples (Fig. S2). According to the theoretical prediction, if divergent selection is sufficiently strong, such that the selection coefficient is higher than the migration rate [49] or the recombination rate [50], GD may occur through the hitchhiking effect. Alternatively, if the combined effect of mild divergent selection is sufficiently strong, then GD may occur for the same reason [51, 52]. This hitchhiking effect was previously coined as genome hitchhiking [39]. We postulate that the observed divergent selection on chromosome 12 and Z chromosomes might be sufficiently strong to contribute to the generation of GD. In a previous study, we also showed that mild divergent selection caused GD between mt-A and mt-B in the FAW population in Mississippi [40]. Here, we hypothesize that the combined effect of mild divergent selection caused GD between mt-A and mt-B in the other geographic populations.

We argue that the mitochondrial COX1 gene[33], and the Z chromosome TPI marker[35] should be used for different purposes. The almost perfect correlation between the genotypes at TPI genes and host-plant groups suggests that the TPI marker can be used to identify host-plant strains. We consider that mitochondrial COX1 is an improper marker to identify *host-plant* strains as we showed that samples with sfR markers have been frequently observed from corns, as also shown in previous studies [36, 40, 42]. The genomic differentiation between mitochondrial subgroups (Fig. 2B) suggests that mitochondrial COX1 can be still used to identify *some* genetic identities within the corn group (i.e., mt-A and mt-B). FAWs from invasive populations are predominantly found in corns [53], and invasive FAWs have sfC-type TPI sequences and sfR-type COX1 sequences [54, 55]. In this case, invasive FAW populations should be considered as sfC, rather than hybrids.

Tessnow et al.[36] showed that allochronic mating patterns may have caused genomic differentiation between sfC and sfR using the samples collected from corn and sorghum, which are preferred host-plants by sfC. They proposed that sfC and sfR should be considered allochronic strains, rather than host-plant strains. However, we believe that this argument is yet to be accepted because they did not analyze samples collected from sfR-preferred host-plants. Interestingly, the differentiation between sfC and sfR appears to be clear when strains were identified from three Z-linked SNVs [56] while the differentiation between sfC and sfR was less clear when mitochondrial COX1 was used. Importantly, because they collected samples in adult stages, the host-plant during larval stages remained unidentified. It is possible that the sfC and sfR identified by Tessnow et al. [36] might correspond to the corn group and the grass groups identified in this study. It is worthwhile to note that Tessnow et al. used different markers to identify strains (i.e., three interspersed SNVs on the Z chromosome. Gene flow from sfR to sfC will increase the relative frequency of grass-fitted alleles (G) to corn-fitted alleles (g) in the corn group. Assortative mating by allochronic mating in the corn group will reduce the efficacy of ecological divergence selection because g-carrying individuals have an increased chance to mate with other individuals with the same strain (sfC or sfR) by assortative mating. Then, the allele frequency of g could be maintained in the sfC despite ecological selection against g, depending upon the relative strength of assortative mating to ecological divergent selection. In other words, the direction of divergence can be different between pre-zygotic and post-zygotic reproductive barriers by recombination, and this unequal direction could interfere with the speciation process. If both preferred host-plants and mating time are determined by the same loci, this interference does not occur and the evolutionary trajectory of differentiation is expected to be the same between differential host-plant adaptation and allochronic mating pattern. If this possibility is true, differential host-plant adaptation and allochronic mating patterns may have additive effects on speciation. The *vrille* gene was proposed to be a gene controlling allochronic mating [25], but we did not find support that this gene caused genetic differentiation between sfC and sfR or mt-A and mt-B (Fig. S6).

We acknowledge that geographic effects on grass-eating FAWs were not taken into account in our analysis because this study is based on a single geographic location for grass-eating FAWs (i.e., grass group). Future studies will need to include more geographic locations both for grass and corn-eating FAW. We also acknowledge that the role of identified genes under divergent selection (Table S4) in speciation is still unclear. If we can narrow down candidate genes in which different alleles generate different fitness in a host-plant species, functional genomic studies could be straightforward to test the role of these candidate genes in host-plant adaptation through RNAi or CRISPR/CAS9 experiment. The resolution of selection scans can be greatly increased when SNVs are phased [57]. Long-read sequencing can be particularly useful for this purpose.

In this study, we posit that host-plant adaptation is one of the main drivers of incipient speciation in the FAW. This speciation process appears to involve divergent selection causing reproductive isolation. The FAW displays differentiated phenotypes potentially causing both prezygotic and postzygotic reproductive barriers. Interestingly, the evolutionary trajectory under these phenotypes may not be uniform in a way of separating the FAW into sfC and sfR. To better understand how interactions between these phenotypes ultimately generated a pattern of genomic differentiation driven by host-plants, future studies should integrate analyses of whole genome sequences from phenotyped individuals collected from a wide range of geographic locations.

## MATERIALS AND METHODS

### Genome assembly

We performed the mapping of available Illumina reads (~80X) [8] from a single sfC individual from a laboratory strain, which was seeded from a population in Guadeloupe in 2000 [8], against an sfC assembly, which was generated from 30X PacBio reads from the same strain in our previous study [40], using SMALT (Sanger Institute). Potential errors in the assemblies were identified using reapr [58]. If an error was found over a gap, the scaffold was broken into two using the same software to remove potential structural errors in the assembly. The broken assemblies were concatenated using SALSA2 [59] or 3D-DNA [60], followed by gap filling with the 80X Illumina reads using SOAP-denovo2 Gap-Closer and with the PacBio reads using LR_GapCloser v1.1 [61].

We observed that 3D-DNA generated a better assembly than SALSA2, as determined by BUSCO analysis (Table S5). Thus, the assembly from 3D-DNA was used in this study. Gene annotation was transferred from the previously generated assemblies (OGS 6.1 on https://bipaa.genouest.org/) to the current assembly using RATT [62].

### Data generation

The samples from Florida were collected from a pasture grass field in Jacksonville (Duvall Co.) and a sweet corn field at Citra (Marion Co.) in Florida (USA) in September 2015 by hand collection. Genomic DNA was extracted from 24 individuals using Dneasy blood and tissues kit, and libraries for Illumina sequencing were generated from 1.0μg DNA for each sample using NEBNext DNA Library Prep Kit with 300bp insertion size. Paired-end genome sequencing was performed using Novaseq S6000 with 150bp reads with 20X coverage for each sample. Adapters in the reads were removed using adapterremoval v2.1.7 [63], followed by mapping the reads against a reference genome with chromosome-sized scaffolds [64], using bowtie2 v2.3.4.1 with −very-sensitive-local preset [65]. Raw Illumina reads from 17 samples from Mississippi (NCBI SRA: PRJNA494340)[8, 40] and 15 samples from Puerto Rico (PRJNA577869) [42] were treated in the same way. Haplotype calling was performed from resulting bam files using GATK v4.1.2.0 [66]. Then, variants were called using GATK v4.1.2.0 [66], and only SNVs were retained. We discarded SNVs if QD is lower than 2.0, FS is higher than 60.0, MQ is lower than 40.0, MQRankSum is lower than - 12.5, or ReadPosRankSum is lower than −8.0. The list of samples is available in Table S6 with detailed information.

### Strain identification

Mitochondrial genomes were assembled and COX1 sequences were extracted using MitoZ [67]. Together with non-FAW COX1 sequences obtained from a previous study [68], a multiple sequence alignment was generated using MUSCLE v3.8.31 [69]. A distance-based phylogenetic tree was reconstructed using FastME v2.1.6 with the F84 evolutionary model [70]. The phylogenetic tree was visualized using iTOL v6[71]. Then, strains were identified from clades containing samples of which strains were identified from previous studies [40, 42].

The strain was also identified using the TPI gene. We extracted a vcf file containing TPI gene from whole the nuclear genomic vcf file using tabix v1.10.2-3 [72]. Principal component analysis was performed using plink v1.9 [73], and two groups according to the strains were identified. Then, the strain of each sample was identified.

### Population genomics analysis

Weir and Cockerham’s F_ST[74]_ was calculated using VCFtools v0.1.15[75]. The window size was 500kb. Statistical genetic differentiation was tested by calculating the proportion of random groups from which the calculated FST is higher than the grouping between the corn and the grass groups or between sfC and sfR in the corn group. DXY in sliding windows was calculated using Dxy [76]. The size of the windows was 500kb and the step size was 100kb. Ancestry coefficient analysis was performed using sNMF v1.2[77]. Selective sweeps were inferred from the composite likelihood of being targeted by selective sweeps from allele frequency spectrums using SweeD v3.2.1 [43]. The grid number per chromosome was 1000. Potential targets of selective sweeps were identified from obvious outliers of composite likelihoods, identified by eyeballing.

Forward simulation was performed using SLiM4[78] to test increased F_ST_ and D_XY_ at divergently selected loci causing reproductive isolation. We chose human conditions to determine the recombination rate (1.19 × 10^-8^)[79], mutation rate (1.2 × 10^-8^)[80], and effective population (3,100) [81]. Simulated populations include two sister populations (Pop A and Pop B) spitted from a common ancestral population. Unidirectional gene flow was allowed from Pop B to Pop A with the migration rate equal to 0.001 to reflect a situation of restricted gene flow from PopB to Pop A by divergent selection in Pop A. Pop A experienced divergent selection with the selection coefficient equal to 0.05. The length of simulated DNA was 2Mb, and divergent selection targeted the middle of sequences. D_XY_ and F_ST_ were calculated from 20kb windows. In total, 50 independent forward simulations were performed and calculated D_XY_ and F_ST_ were averaged.

## LIST OF ABBREVIATIONS

COX1: Cytochrome *c* oxidase subunit 1
FAW: Fall Armyworm
GD: Genomic differentiation
PCA: principal component analysis
sfC: *Spodoptera frugiperda*, corn strain
sfR: *Spodoptera frugiperda*, rice strain
SNP: Single nucleotide polymorphism
TPI: Triosephosphate Isomerase
USA: United States of America

## DECLARATIONS

### Ethics approval and consent to participate

Not relevant.

### Guidelines and regulations in methods

Not relevant.

### Consent for publication

Not relevant.

### Availability of data and materials

The raw reads of these samples are available from NCBI SRA (PRJNA639296). The reference genome assembly used in this study (ver7) is available at BIPAA (https://bipaa.genouest.org/sp/spodoptera_frugiperda_pub/download). Computer programming scripts used in this study are available on request.

### Authors’ contributions

EF prepared the resequencing dataset and generated Fig. 1. KD contributed to generating Fig. 3 and 4.SG and EA contributed to preparing the resequencing dataset. FL generated the reference genome assembly. RLM, GJK, and NN provided samples used in this study. KN generated Fig. 2, 4, and 5. KN conceived and designed the analyses. All authors participated in writing the paper.

## Acknowledgments

We are grateful to the genotoul bioinformatics platform Toulouse Occitanie (Bioinfo Genotoul, https://doi.org/10.15454/1.5572369328961167E12), GenOuest (https://www.genouest.org/), and BioInformatics Platform for Agroecosystem Arthropods (https://bipaa.genouest.org/is/) for providing help and/or computing and/or storage resources.

