## Supplementary figures and tables for "Host-plant adaptation as a driver of incipient speciation in the fall armyworm (*Spodoptera frugiperda*)"

**Table S1**. Summary statistics of genome assemblies showing contiguity and correctness of the assemblies (BUSCO) generated in this study and other published assemblies.

|  | Statistics | Current assembly | Gimenez *et al*.[1] | Nam *et al.*[2] | Gouin *et al.*[3] | | Liu *et al.*[4] | |
| --- | --- | --- | --- | --- | --- | --- | --- | --- |
|  |  |  |  |  | sfC | sfR | Male | Female |
| contiguity | Assembly size | 385,049,154 | 384,455,365 | 379,902,278 | 437,873,304 | 371,020,040 | 543,659,128 | 531,931,622 |
|  | number of sequences | 495 | 125 | 1,054 | 41,577 | 29,127 | 21,840 | 27,258 |
|  | N50 (bp) | 13,567,047 | 13,151,234 | 1,129,192 | 52,781 | 28,526 | 14,162,803 | 13,967,093 |
|  | L50 | 13 | 13 | 91 | 1,616 | 3,761 | 16 | 17 |
|  | N90 (bp) | 10,609,592 | 8,473,354 | 165,330 | 3,545 | 6,422 | 6,440 | 5,122 |
|  | L90 | 26 | 27 | 421 | 18,789 | 13,881 | 3,030 | 5,612 |
| BUSCO | Complete | 1,601 | 1,601 | 1,616 | 1,461 | 1,551 | 1,576 | 1,577 |
|  | Complete and single-copy | 1,577 | 1,573 | 1,573 | 1,276 | 1,518 | 1,442 | 1,480 |
|  | Complete and duplicated | 25 | 28 | 43 | 185 | 33 | 134 | 97 |
|  | Fragmented | 19 | 20 | 11 | 127 | 69 | 45 | 48 |
|  | Missing | 37 | 37 | 31 | 70 | 38 | 37 | 33 |
|  | Total | 1,658 | 1,658 | 1,658 | 1658 | 1,658 | 1,658 | 1,658 |

**Table S2**. The information of each sample used in this study.

| ID | location | host-plant | TPI | mt-COX1 |
| --- | --- | --- | --- | --- |
| FCC1 | Florida | Corn field | sfC | sfC |
| FCC2 |  |  | sfC | sfC |
| FCC3 |  |  | sfC | sfC |
| FCC4 |  |  | sfC | sfC |
| FCC5 |  |  | sfC | sfC |
| FCC6 |  |  | sfC | sfC |
| FCC7 |  |  | sfC | sfC |
| FCC8 |  |  | sfC | sfC |
| FL.16 |  |  | sfC | sfC |
| FL.17 |  |  | sfC | sfC |
| FL.18 |  |  | sfC | sfC |
| FL.19 |  |  | sfC | sfC |
| FL.20 |  |  | sfC | sfC |
| FGJ10 |  | pasture grass | sfR | sfR |
| FGJ11 |  |  | sfR | sfR |
| FGJ12 |  |  | sfR | sfR |
| FGJ2 |  |  | sfR | sfR |
| FGJ3 |  |  | sfR | sfR |
| FGJ4 |  |  | sfR | sfC |
| FGJ5 |  |  | sfR | sfR |
| FGJ6 |  |  | sfR | sfR |
| FGJ7 |  |  | sfR | sfR |
| FGJ8 |  |  | sfR | sfR |
| FGJ9 |  |  | sfR | sfR |
| MS_C_C1 | Mississippi | Corn field | sfC | sfC |
| MS_C_C2 |  |  | sfC | sfC |
| MS_C_C3 |  |  | sfC | sfC |
| MS_C_C4 |  |  | sfC | sfC |
| MS_C_C5 |  |  | sfC | sfC |
| MS_C_C6 |  |  | sfC | sfC |
| MS_C_C7 |  |  | sfC | sfC |
| MS_C_C8 |  |  | sfC | sfC |
| MS_C_C9 |  |  | sfC | sfC |
| MS_R_R2 |  |  | sfC | sfR |
| MS_R_R3 |  |  | sfC | sfR |
| MS_R_R4 |  |  | sfC | sfR |
| MS_R_R5 |  |  | sfC | sfR |
| MS_R_R6 |  |  | sfR | sfR |
| MS_R_R7 |  |  | sfC | sfR |
| MS_R_R8 |  |  | sfC | sfR |
| MS_R_R9 |  |  | sfC | sfR |
| PR1 | Puerto Rico | Corn field | sfC | sfC |
| PR12 |  |  | sfC | sfC |
| PR14 |  |  | sfC | sfR |
| PR15 |  |  | sfC | sfC |
| PR16 |  |  | sfC | sfC |
| PR18 |  |  | sfC | sfC |
| PR19 |  |  | sfC | sfC |
| PR27 |  |  | sfC | sfR |
| PR29 |  |  | sfC | sfC |
| PR30 |  |  | sfC | sfR |
| PR31 |  |  | sfC | sfC |
| PR32 |  |  | sfC | sfC |
| PR33 |  |  | sfC | sfC |
| PR35 |  |  | sfC | sfR |
| PR5 |  |  | sfC | sfC |

**Table S3**. Position of targets of selective sweeps specific to the corn or grass group

| Group under selective sweep | Chromosome | Start | End |
| --- | --- | --- | --- |
| Grass group | 12 | 17,246,524 | 17,452,113 |
| Corn group | Z | 15,849,810 | 16,376,377 |
| Corn group | Z | 18,693,271 | 19,177,712 |

**Table S5**. Summary statistics of genome assemblies showing contiguity and correctness of the assemblies generated using SALSA2 and 3D-DNA.

|  |  | SALSA2 | 3D-DNA |
| --- | --- | --- | --- |
| Contiguity | Number of scaffolds | 390 | 495 |
|  | Size | 384Mb | 385Mb |
|  | N50 | 22.8Mb | 13.6Mb |
|  | L50 | 6 | 13 |
|  | N90 | 0.8MB | 10.6Mb |
|  | L90 | 30 | 26 |
| Correctness  (BUSCO) | Complete | 1,601 | 1,601 |
|  | Complete and single-copy | 1,570 | 1,577 |
|  | Complete and duplicated | 31 | 25 |
|  | Fragmented | 22 | 19 |
|  | Missing | 35 | 37 |

**Table S6**. The list of samples used in this study.

| State in the USA | Village | Host plant | Sampling method | Sampling year | The number of samples | Source |
| --- | --- | --- | --- | --- | --- | --- |
| Puerto Rico | Santa Isabel | Corn | handpick | 2009 | 15 | [1] |
| Mississippi | Stoneville | Corn | handpick | 2009 | 17 | [2, 3] |
| Florida | Citra | Corn | handpick | 2015 | 13 | This study |
| Florida | Jacksonville | Grass | handpick | 2015 | 11 | This study |

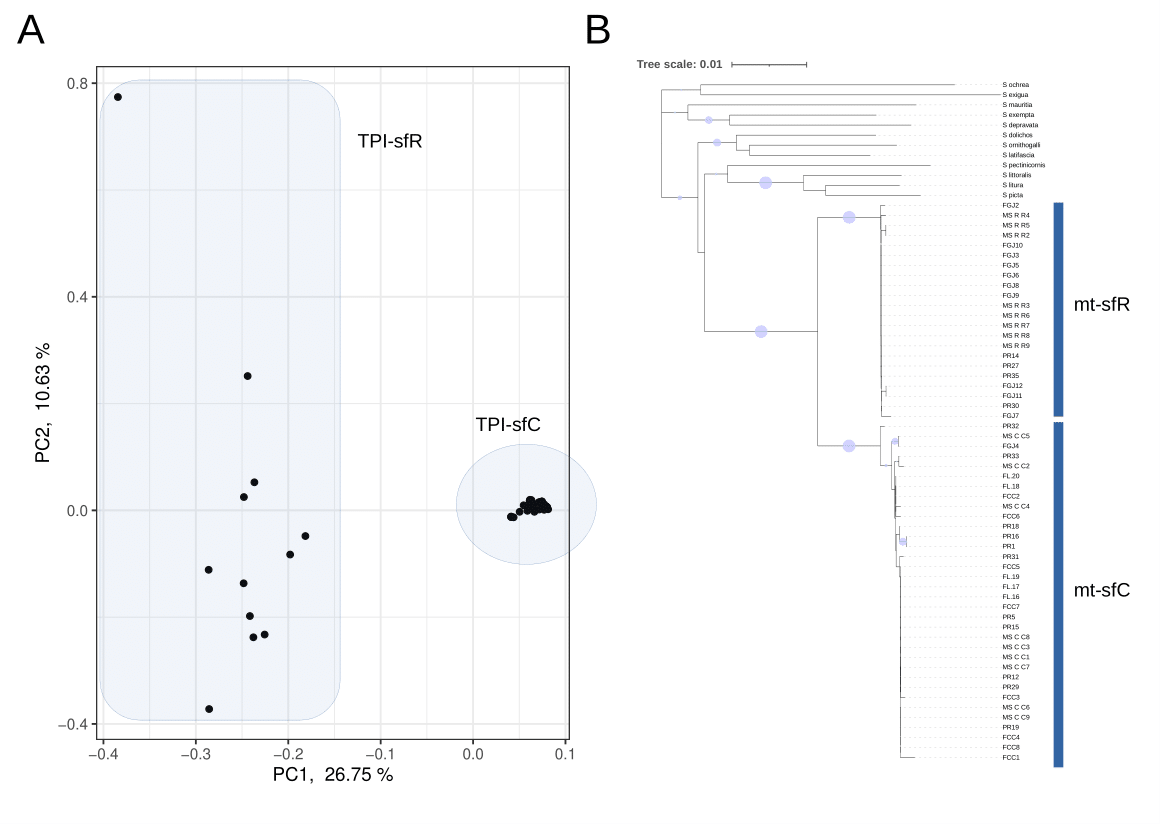
**Figure S1**. Identification of strains. A. Principal component analysis from TPI genes. B. Phylogenetic tree from mitochondrial COX1 marker gene. Non-parametric bootstrapping values above 70% were shown with circles on the branches. Rooting was performed based on the reported phylogenetic tree in *Spodoptera*[5].

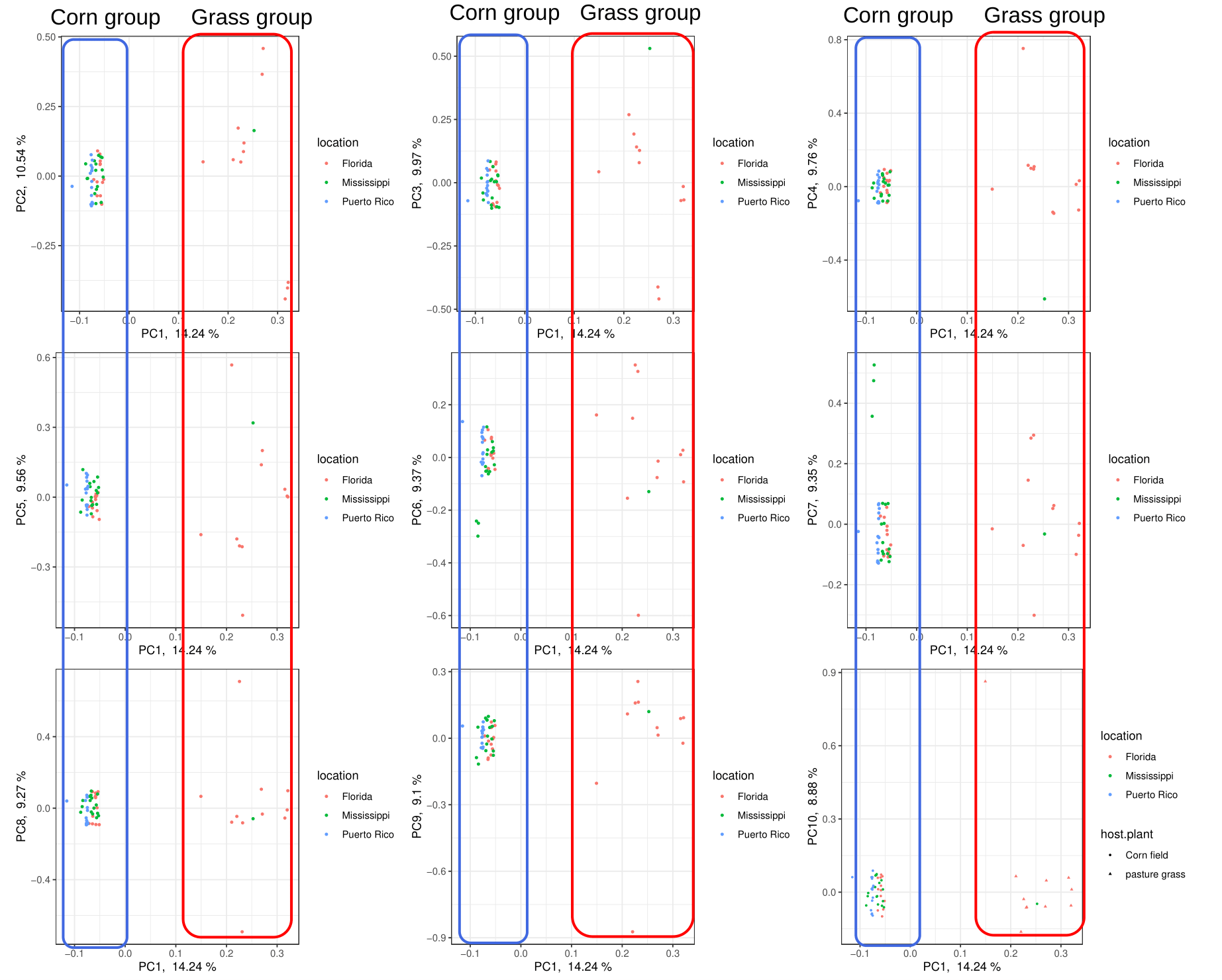

**Figure S2**.The result of the principal component analysis from whole nuclear genome sequences with the information of sampling locations. The grouping according to geographic populations was not observed from the first principal component to the tenth principal component. Please note that geom_jitter(height=.1,width=0) was used on ggplot2 in R for visualization purpose.

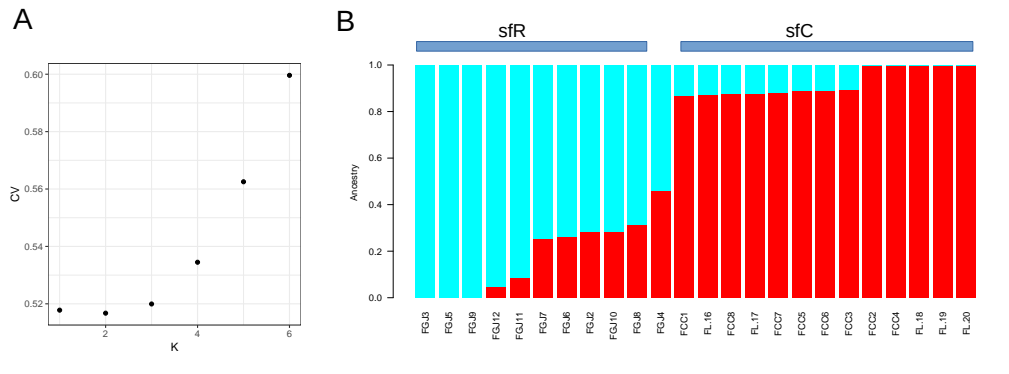
**Figure S3**. Ancestry coefficient analysis using the samples from Florida. A. The cross-validation score was lowest when K =2. Therefore, we chose the result of ancestry coefficient analysis at K = 2. B. sfC and sfR samples exhibited differentiated ancestry. FG4 showed almost the same proportion of sfC and sfR ancestries.

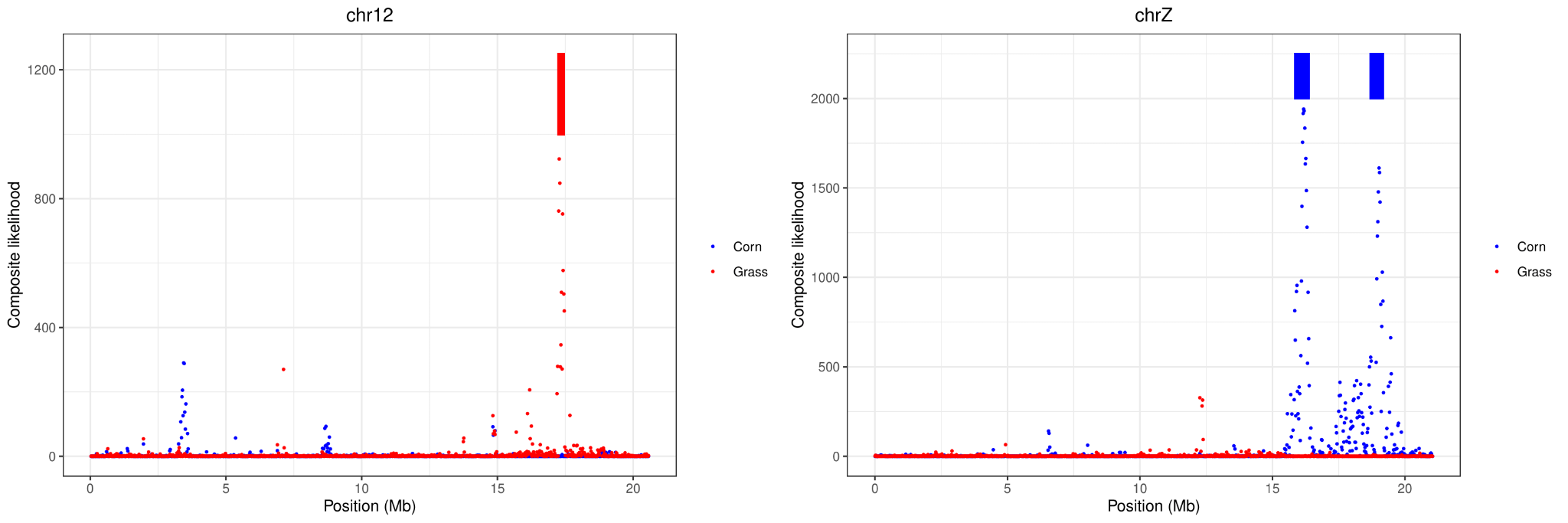
**Figure S4**. The composite likelihood of being targeted by selection on chromosome 12 and Z chromosomes. The red and blue bars represent the outliers in the grass and corn groups, respectively.

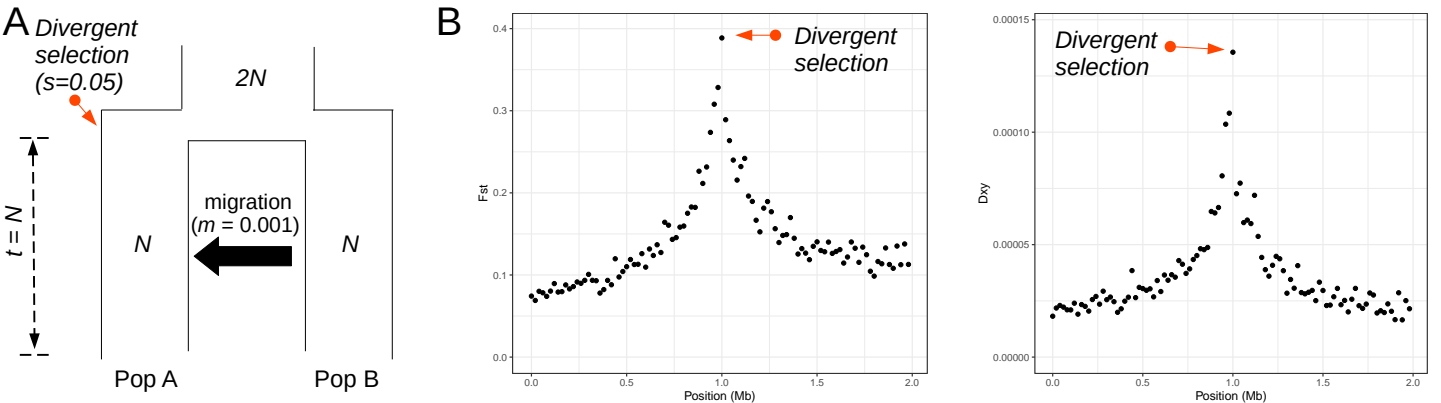
**Figure S5.** **Forward simulation to test increased D_XY_ and F_ST_ at divergently selected loci causing reproductive isolation** A. Model used in the simulation. Ancestral population with an effective population equal to 2*N* split into two populations (Pop A and Pop B) of which the population size was *N*, *N* generations ago. Unidirectional migration existed from Pop B and Pop A. The rate of migration (*m*) was 0.0001 per generation. Pop A experienced divergent selection at the beginning of the split into Pop A and Pop B. Selection coefficient (*s*) was 0.05. *N* equals to human effective population size, 3,100. B. F_ST_ and D_XY_ along the simulated sequences. The red arrow bars indicate the divergently selected position.

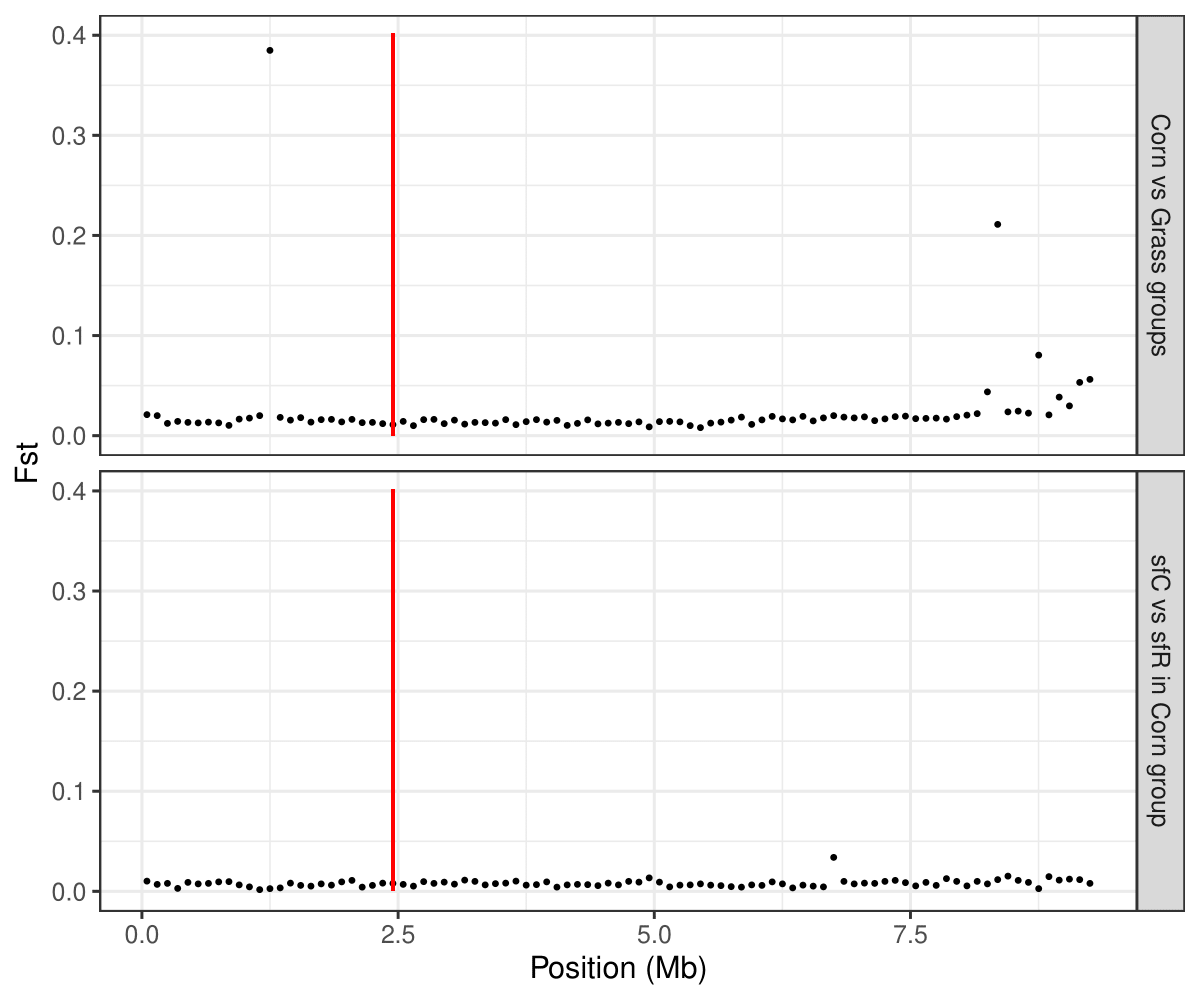
Figure S6. F_ST_ between corn and grass groups or between sfC and sfR in the corn group was calculated in 100kb window along chromosome 23. The red vertical bar indicates a locus of the *vrille* gene.
